# Extensive adaptive changes in bat interferon pathway reveal specific molecular functions at the forefront of host–virus coevolution

**DOI:** 10.1101/2025.09.17.676773

**Authors:** Stéphanie Jacquet, Adil El Filali, Lucie Etienne, Dominique Pontier

## Abstract

The interferon (IFN) response is a fundamental component of the mammalian innate immune system, orchestrating a broad antiviral state through a network of interferon-stimulated genes (ISGs). In bats, which are natural reservoirs for diverse viruses, several ISGs exhibit lineage-specific evolutionary adaptations, reflecting unique immune strategies. While previous research has typically focused on individual ISGs or broad-scale genomic analyses, here, we conducted an evolutionary analysis of type I IFN-upregulated genes across diverse bat species to elucidate the forces shaping the full repertoire of their ISG responses. Our results reveal that key components of bats antiviral signaling, including viral dsRNA sensors, PARP-mediated ADP-ribosylation enzymes, and cytokine/chemokine signaling molecules, have been recurrent targets of strong positive selection. Notably, immune sensors such as TLR3, RIG-I, and MDA5 show adaptive changes in their RNA-binding domains, suggesting modified viral recognition. Members of the PARP family exhibit repeated selection in both macro and catalytic domains, consistent with intense host–virus conflict and immune modulation. Strikingly, chemokine genes—especially CXCL10 and CXCL16—and their receptors display some of the strongest and most concerted selection signatures, particularly in ligand-binding loops, consistent with arms-race coevolution potentially driven by viral mimicry. Several positively selected genes have pleiotropic functions in tissue repair, inflammation, and tumor control, suggesting that some of these adaptations may influence several physiological pathways. Altogether, our findings uncover a bat-specific antiviral architecture shaped by concerted adaptive evolution, highlighting a balance between viral control and immune tolerance, likely underlying bats’ exceptional resilience to disease.

**Author summary:** Bats exhibit a remarkable capacity to host a wide range of viral families, including viruses highly pathogenic to humans, while rarely displaying overt clinical symptoms. This resilience suggests unique immune adaptations to control infections while avoiding harmful inflammation. To investigate the genetic basis of this trait, we conducted a comparative evolutionary analysis of genes involved in the type I interferon (IFN) pathway, a core component of the mammalian antiviral response, across multiple bat species and other mammals. Our results reveal strong signatures of positive selection on key antiviral effectors in bats, including sensors of viral nucleic acids, ADP-ribosylating PARP enzymes, and components of chemokine signaling pathways. Notably, several chemokines and their receptors display accelerated evolution, pointing to a critical role in modulating immune responses. We further identify viral chemokine-binding proteins (vCKBPs) from poxviruses as one likely driver of natural selection, suggesting ongoing evolutionary arms races between host immune signaling and viral immune evasion. Overall, these findings fit with a model in which bats have evolved a balance with potent antiviral activity and controlled immune activation. Elucidating these mechanisms offers promising insights for developing novel approaches to managing inflammation and viral infections in humans.

## Introduction

As a central component of cell-autonomous defense, the interferon (IFN) pathway plays a key role in mammalian innate immunity^1^. Upon viral infection, host cells detect pathogen-associated molecules, activities, or stress signals *via* specialized sensors, including pattern recognition receptors (PRRs), thereby triggering the production of cytokines, including IFNs. These IFNs bind to specific cell surface receptors, activating intracellular signaling cascades such as the Janus kinase-signal transducer and activator of transcription (JAK–STAT) pathway. This signaling induces the transcription of hundreds of interferon-stimulated genes (ISGs), several of which act as key antiviral effectors by restricting viral replication within infected cells and enhancing resistance in neighboring cells^1^.

The central role of IFN signaling in antiviral defense led to sustained efforts to characterize the functional repertoire of this pathway across mammals. More recently, this interest has extended to bats, which are increasingly recognized for their remarkable ability to balance viral tolerance with effective immune resistance^2,3^. As the second most diverse mammalian order, bats host a broad spectrum of viral pathogens, including viruses that are closely related to major zoonotic viruses such as SARS-related coronaviruses and henipaviruses^4–6^. Despite this viral diversity, many bat species appear to remain asymptomatic upon infection, in contrast to the severe pathogenicity observed in other mammals^5^. This feature has fueled investigations into how bats control viral infections without triggering excessive inflammation. Growing evidence suggests that bats have evolved immune adaptations that dampen inflammation and enhance tolerance, including genetic changes in natural killer cell receptors^7^, inflammasome components (e.g., loss of PYHIN and AIM2 family genes^8,9^), and signaling proteins (e.g., STING^10^, ASC2^11^, and IRF3^12^). These adaptations are counterbalanced by mechanisms that enhance antiviral resistance, potentially enabling effective viral control without triggering damaging immune responses. Transcriptomic studies have shown that some pteropodid species constitutively express IFN-α and downstream ISGs even in the absence of viral infection^13^. In contrast, other pteropodid species such as *Rousettus aegyptiacus* exhibit high baseline expression of an expanded repertoire of IFN-ω genes, but no detectable constitutive IFN-α activity^7^. These differences extend to the Yangochiroptera suborder, where species like *Myotis* lack several ISGs found in pteropodids^14^, highlighting the remarkable species-specific diversity in IFN regulation.

Beyond these distinct expression patterns of IFN signaling, several canonical ISGs with broad antiviral activities, such as PKR, APOBEC, BST2, ISG15 or RTP4, exhibit strong and/or species-specific signature of adaptation in bats^15–18^. This is supported by comparative analyses of high-quality bat genomes and functional experiments, which have revealed an excess of immune gene adaptations both along the ancestral branch of bats and across multiple bat lineages^18–21^. However, despite these valuable insights, most studies have primarily focused either on individual ISGs or on genome-wide patterns, missing some key rapidly evolving ISGs or their complex evolutionary history, respectively. As a result, the evolutionary dynamics of the full IFN-upregulated gene repertoire across the chiropteran radiation remains only partially explored.

To address this gap, we took advantage of publicly available transcriptomic data from type I IFN-stimulated *Pteropus* and *Myotis* bat cell lines^22^ to investigate the evolutionary diversification of ISGs in bats. We first showed that nearly half of the bat ISGs studied exhibited strong signatures of positive selection, consistent with recurrent or ancient selective pressures, possibly reflecting both host–virus arms races and broader physiological adaptations linked to bat-specific traits such as flight. Notably, genes involved in double-stranded RNA binding, cytokine activity, and poly-ADP ribosyltransferase functions emerged as recurrent targets of adaptive changes. We then focused on the cytokine and chemokine gene families, conducting detailed evolutionary analyses that revealed trajectories potentially linked to bats’ unique capacity to mount effective antiviral response while maintaining immune homeostasis to viral threats.

## Results

### 1. Strong diversifying selection has shaped almost half of the bat type I interferome

To determine how bat ISGs have evolved during chiropteran diversification, we performed interferome-wide evolutionary analyses using publicly available data from Shaw et al. 2017^22^ (https://isg.data.cvr.ac.uk/), which experimentally compared IFN-stimulated transcriptomes from bats (*Pteropus vampyrus* and *Myotis lucifugus*), human, brown rat, cattle, pig, sheep, and horse cell lines. Using a cut-off of Log_2_ fold-change > 2 and a false discovery rate (FDR) < 0.05 for upregulated genes, we identified 327 genes as part of the bat interferome (Table S1). After excluding 16 genes (families) due to lack of annotation, misidentification, or excessive duplications, 311 genes were retained for analyses. Interestingly, while approximately half of these genes (n = 166) were upregulated in both bat species, one-third were specific to *P. vampyrus* (n = 112) and the remaining (n = 33) were specific to *M. lucifugus* (Fig. 1A), highlighting marked interspecific and/or inter-cell type differences in bat IFN responses.

**Figure 1.**
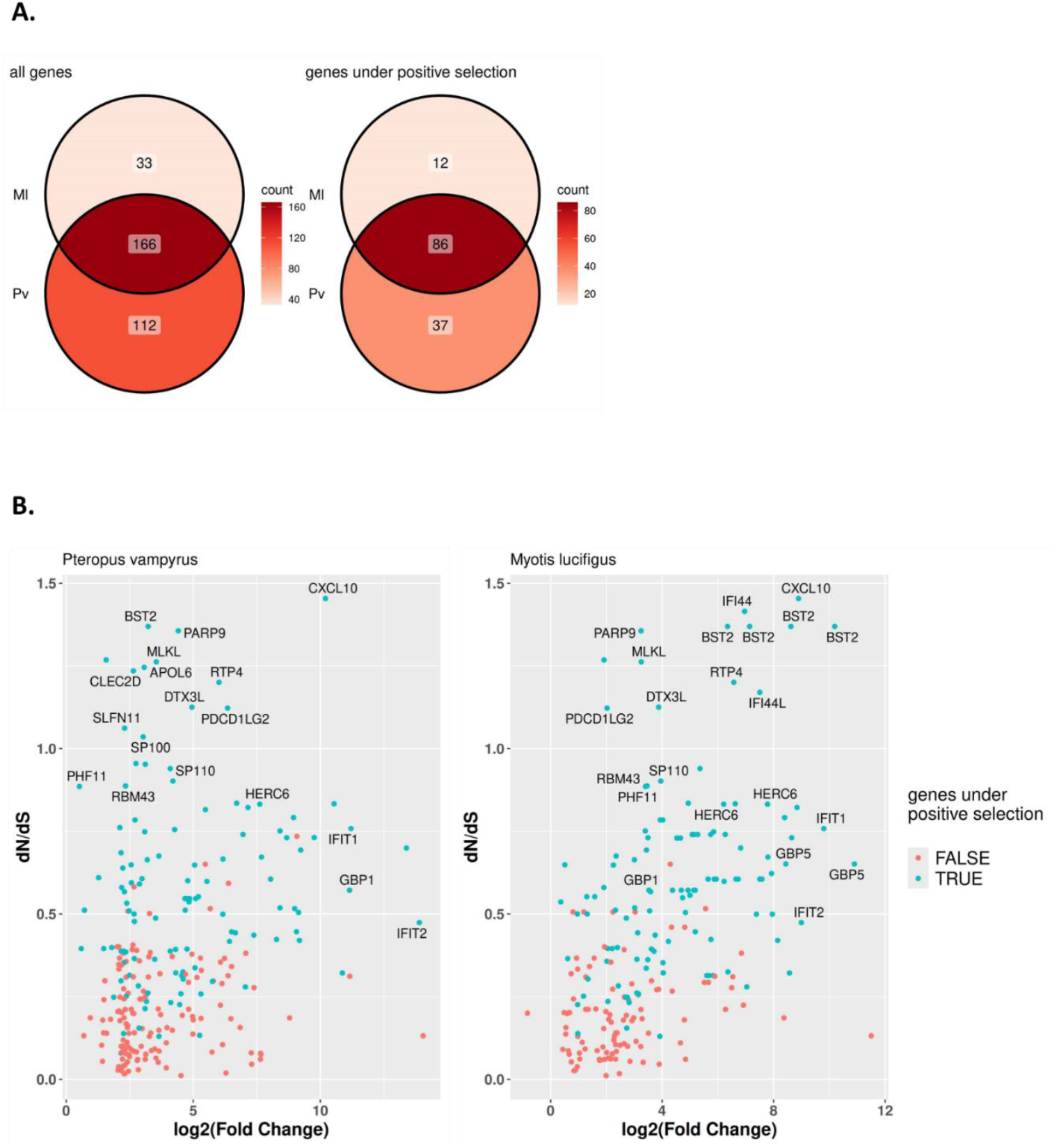
Adaptive evolution in bats ISGs. (A) Venn diagram showing the number of ISGs (with a Log_2_(fold change) > 2) identified in *Pteropus vampyrus* (Pv) and *Myotis lucifigus* (Ml) 22, which were screened for signatures of positive selection. (B)Average dN/dS values estimated using the CodemL M8 model plotted against log_2_(fold change) in *P. vampyrus* and *M. lucifugus*, with ISGs under significant positive selection (p-value < 0.05) highlighted in blue. IFN-stimulated upregulation is significantly correlated to the average dN/dS (Spearman’s rank correlation, p-value = 2.7 10^-06^)

We therefore screened these 311 ISGs to detect signatures of positive selection (i.e., elevated ratio of non-synonymous to synonymous substitution rates; dN/dS > 1). To ensure robust inference, we employed three complementary methods: the Branch-site Unrestricted Statistical Test for Episodic Diversification (BUSTED)^23^ implemented in HYPHY^24^, along with two likelihood ratio tests from the Codeml package of PAML (model comparisons M1 vs. M2 and M7 vs. M8). A gene was considered to be under positive selection if at least two of the three methods yielded statistically significant results. Despite this conservative approach, nearly half of the tested genes (n = 139) showed evidence of positive selection (LRT p-value < 0.05 for at least two methods), many of which are potent mammalian immune effectors, including cGAS, OASs, SAMHD1, SLFN11, SHFL, TRIM5, TRIM22, or ZAP. Plotting average dN/dS ratios against IFN Log_2_(FC) for each gene revealed a significant positive correlation between IFN-stimulated upregulation and the strength of selective pressure (Spearman’s rank correlation, p-value = 2.7 10^-06^, Fig. 1b, Supplementary Fig. 1A). Importantly, among the top IFN-upregulated and positively selected genes, we identified: (i) well-documented bat antiviral effectors: BST2, GBP1, GBP5, RTP4 and RIPK3/MLKL axis^16,17,25–29^; (ii) important antiviral effectors extensively studied in primates but still underexplored in bats: CXCL10, DTX3L, IFI44/44L, IFIT1-2, PARP9, SLFN11, TRIM22; and (iii) novel or poorly characterized proteins as immune effectors, APOL6, CLEC2D, PDCD1LG2 (PD-L2), PHF11, RBM43, SP100 (Fig. 1B), which we propose as strong candidates for future experimental studies in bats’ innate immunity. To start exploring the latter, we extended our evolutionary analyses to PHF11 and RBM43 orthologs in primates and rodents. Interestingly, both proteins also showed strong signatures of positive selection (p-value < 10^-6^) across these mammalian orders, suggesting that these fast-evolving proteins are potentially overlooked components in the global mammalian IFN response.

### 2. dsRNA sensing, Poly-ADP ribosyltransferase, and cytokine activities are among the most rapidly evolving functions in the bat type I interferon pathway

To identify which pathways and molecular functions have been enriched in adaptive selection in bats, we performed Gene Ontology (GO) search on the positively selected genes, using the entire ISG dataset as background. We found a significant enrichment of positively selected genes across various molecular functions involved in both early and late phases of the innate immune responses (Fig. 2A). Notably, the highest enrichment of positively selected genes was observed for dsRNA binding, poly-ADP ribosyltransferase, and cytokine/chemokine activities, which primarily function in the early and intermediate phases of the innate immune response, and share overlapping functional roles and evolutionary pressures (Fig. 2B-C).

**Figure 2.**
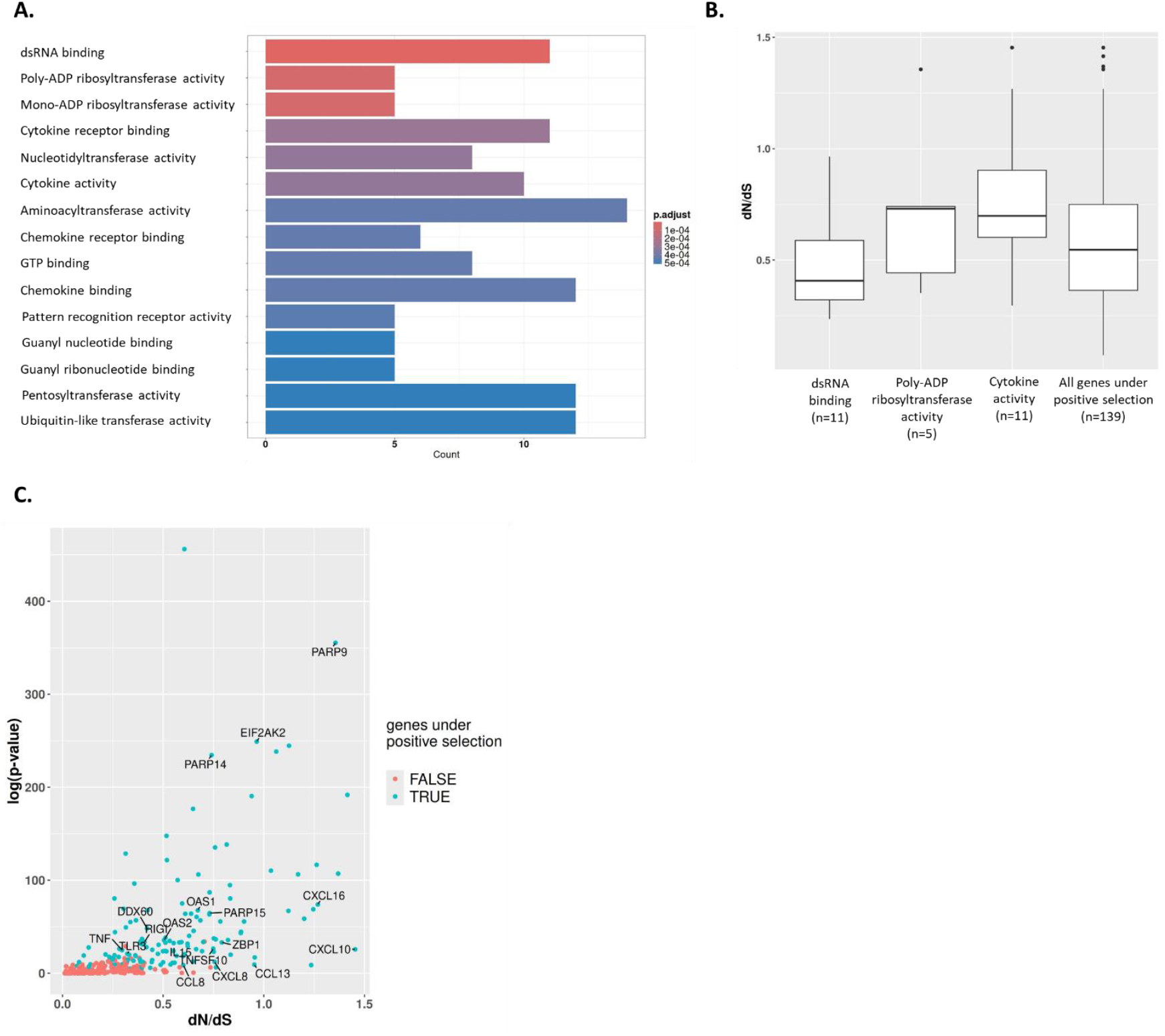
dsRNA sensing and PARP ADP-ribosyltransferase activities are major targets of positive selection in bats. (A) Molecular enrichment analysis highlighting key functions under adaptive selection. (B) Average dN/dS for the four most enriched molecular functions among the ISGs under significant positive selection (C) Log-transformed p-values from Codeml model comparisons (M7 vs. M8), indicating the strength of positive selection acting on genes associated with these enriched molecular functions.

During the early phases of infection, viral dsRNA is sensed by pattern-recognition receptors (PRRs), including Toll-like receptors (TLRs), RIG-I-like receptors (RLRs), and antiviral effectors such as PKR and OAS1, which trigger or enhance the expression of interferon-stimulated genes (ISGs) and pro-inflammatory cytokines. Here, the GO terms associated with dsRNA sensing activity include well documented PRRs like MDA5, RIG-I, and TLR3, along with potent and key antiviral effectors like DDX60, EIF2AK2/PKR, OASs, ZBP1, and ZNFX1. Notably, these dsRNA sensors exhibited strong signatures of positive selection, with significant excess of non-synonymous substitutions (Fig. 3A), mostly concentrated in their dsRNA binding regions (Fig. 3A, B). This pattern, which differs from other mammalian orders^e.g.15,30–32^, suggests a potential modulation of ligand recognition or binding affinity to viral dsRNA, possibly impacting antiviral sensing mechanisms.

**Figure 3.**
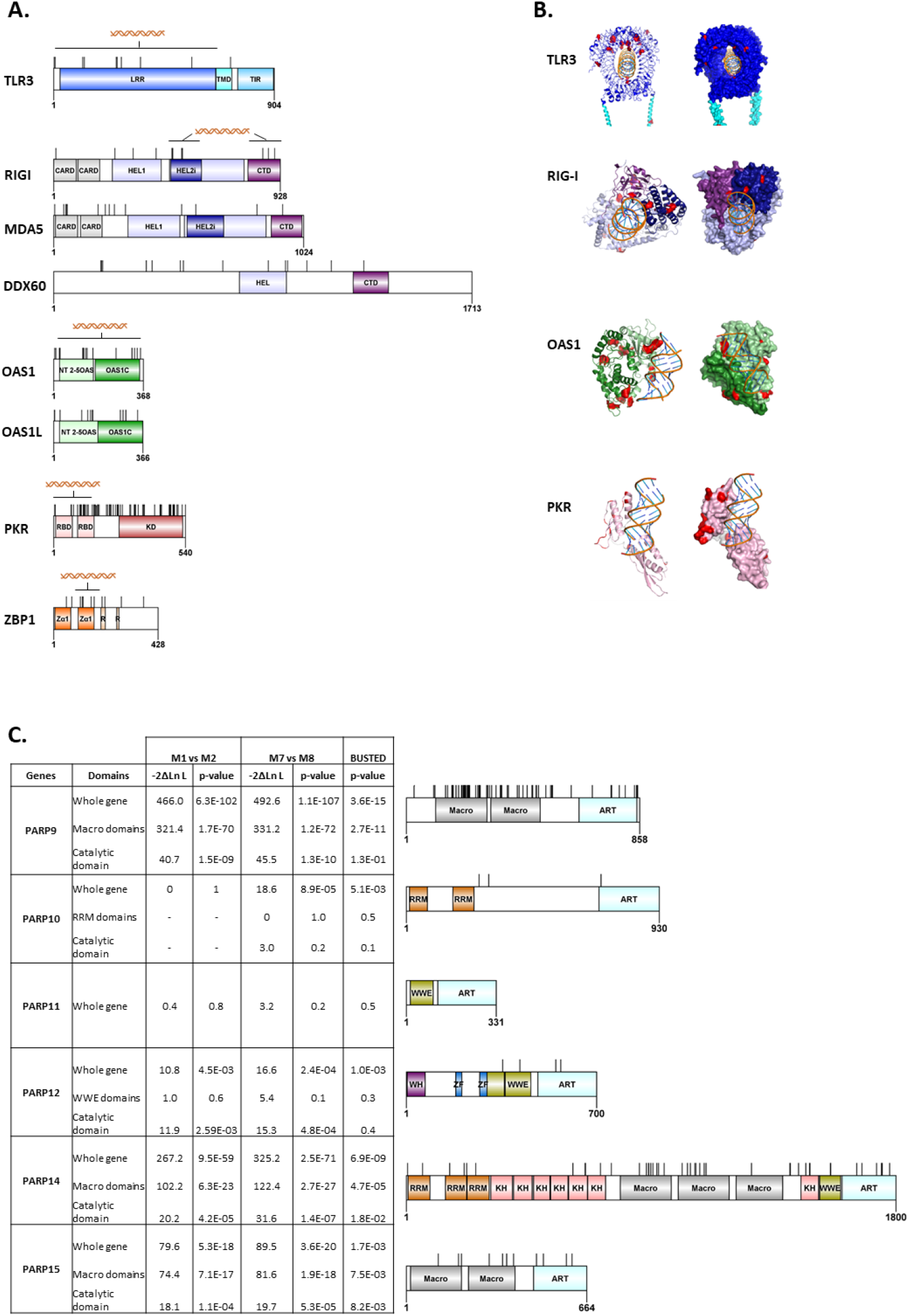
dsRNA binding domains and PARP macro domains exhibit strong signatures of adaptive selection. (A) Protein 2D diagrams showing positively selected sites (BEB > 0.95 for Codeml models; P > 0.95 for FUBAR; p-value < 0.05 for FEL and MEME) along key domains, including hotspots located in the dsRNA binding domains of several bat sensors. (B) 3D protein-dsRNA docking models inferred with HDOCK. Positively selected sites are shown in red within the dsRNA binding domains. PARP evolution in bats. Results of positive selection analyses with 2D protein diagrams mapping positively selected sites onto PARP proteins and domains. *Pteropus giganteus* was used as the reference species for site positions and protein structure models.

A second major target of positive selection was the family of Poly(ADP-ribose) polymerases (PARPs), which are involved in diverse cellular processes^33,34^. In the context of innate immunity, several PARPs regulate antiviral responses by modulating interferon signaling, inflammasome activation, or viral replication. These proteins function by transferring ADP-ribose from NAD^+^ to target proteins, thereby altering their function or stability. In the studied bat species, six PARP members (PARP9 to PARP15), known as important antiviral restriction factors in primates, are highly upregulated upon IFN treatment (Log_2_(FC) > 4, Table S1). All, except PARP11, showed evidence of positive selection (Fig. 3C), pinpointing their potential role as crucial antiviral effectors in bats. Notably, PARP9, 14, and 15 were among the fastest-evolving PARP members analyzed and exhibited highly localized (i.e. site-specific) adaptation signatures in their macro and catalytic domains (Fig. 3C), which could induce a species-specific antiviral response in bats upon viral infections.

Finally, we found that genes involved in cytokine activity, including IL15, and members of the TNF superfamily (TNFα, TNFSF10, TNFSF13B, TNFRSF14) have been targets of diversifying selection (Supplementary Fig. 2). Notably, members of chemokine gene family (CCL8, CCL13, CCL20, CXCL2, CXCL8, CXCL10, CXCL16), which play a critical role at the interface between innate and adaptive immunity, exhibited some of the highest average dN/dS ratios and rank among the most strongly upregulated gene categories in this dataset (Fig. 1B, 2B-C).

### 3. Concerted evolution of bat chemokines shows adaptive signatures typical of host-virus conflicts and ligand-receptor co-adaptation

Given the evolutionary and functional relevance of cytokines and chemokines, we characterized these genes in further details. Cytokines are small proteins that mediate communication between immune cells and regulate immune activation, inflammation, and tissue repair. They include growth factors, interferons (IFNs), tumor necrosis factor (TNF), interleukins (ILs), and chemokines. The latter are a subset of cytokines that act through G-protein-coupled receptors and play a key role in innate immunity, by guiding immune effector cells to sites of infection or inflammation and coordinating immune cell interactions^35^. Based on the number and arrangement of conserved cysteines in their amino acid sequences, chemokines are classified into four subfamilies: CXC, CC, XC, and CX3C.

Detailed evolutionary analyses of IFN-upregulated chemokines revealed extensive adaptive changes in seven of the nine genes tested, both at the gene and codon levels, particularly in CCL20, CXCL2, CXCL8, CXCL10 and CXCL16 (Table S1, p-values < 0.05). However, due to the extensive duplication of CXCL2 (also known as the GRO protein) in bats, with up to three undistinguishable copies in some species (Supplementary Fig. 3), we excluded it from further analyses to avoid bias in selection analyses. Applying the aBSREL model to a concatenated chemokine alignment across mammals, we detected strong signals of episodic positive selection predominantly in bat lineages, particularly within the Mormoopid and Vespertilionid lineages (Fig. 4A). This pattern was consistently observed in gene-by-gene analyses restricted to bats (supplementary Fig. 4). These findings highlight bats as key targets of chemokine adaptation among mammals. Moreover, average dN/dS ratios in these bat chemokines exceeded those in other mammals, particularly CCL13, CXCL8, CXCL10, and CXCL16 (Fig. 4C). Among these, CXCL16 also exhibited adaptive changes in three of the four tested mammalian orders, highlighting its broad involvement in genetic conflicts (Supplementary Fig. 4).

**Figure 4.**
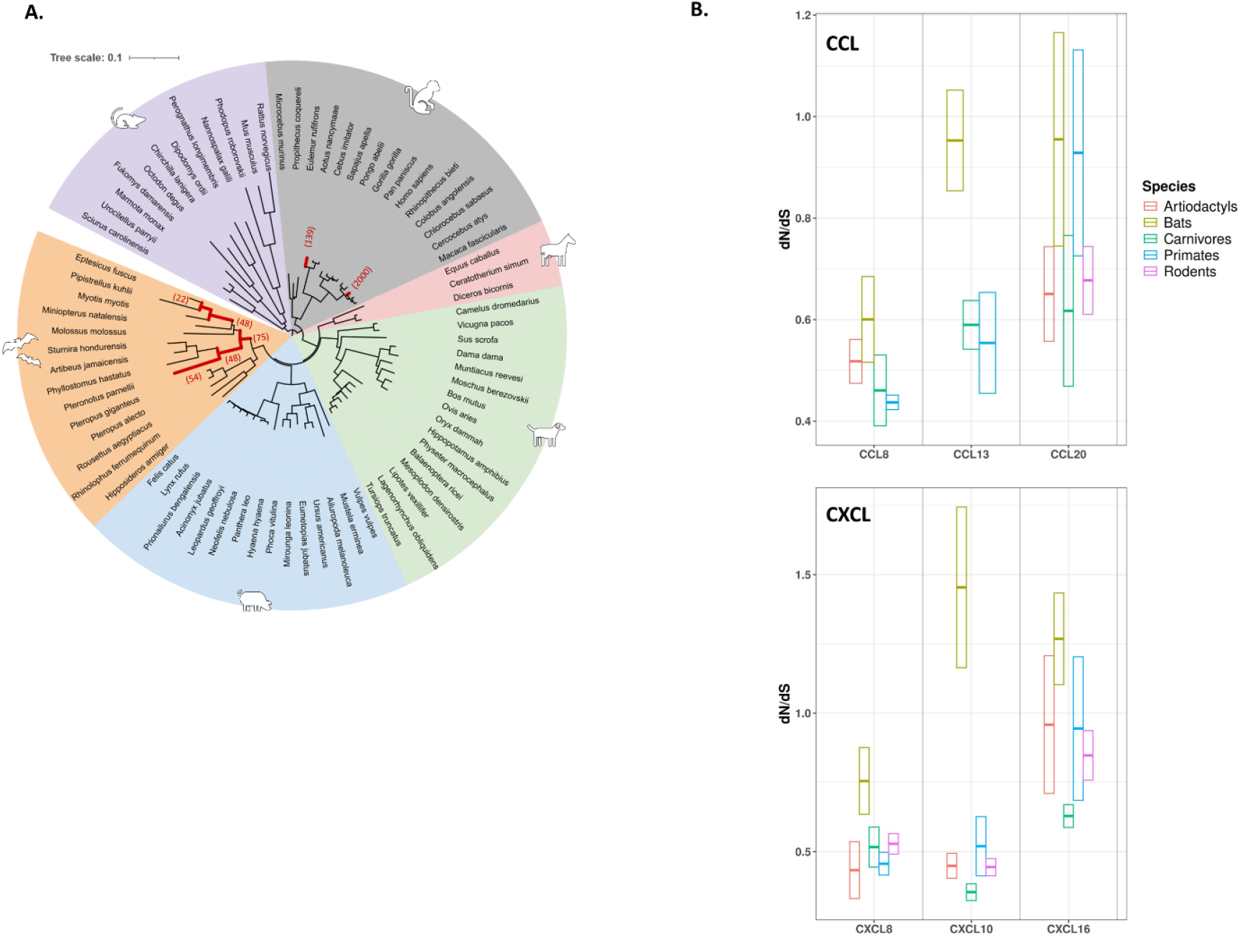
Bats are major targets of chemokine adaptation among mammals. Maximum likelihood phylogenetic trees of (A) a concatenated mammalian dataset of chemokines (CCL5, 8, 13, 20, and CXCL8, 10, 11, 16) and (B) CXCL16 alone, showing branches under significant positive selection (p-value <0.05, in red). Numbers in brackets indicate the estimated values of the ω (dN/dS) value for each branch. The scale bar indicates the proportion of genetic variation. (C) Average dN/dS ratios for mammalian chemokines.

All rapidly evolving bat chemokines exhibited a common evolutionary pattern: positive selection was consistently concentrated within the CC or CXC motif domains encompassing β-strands (Fig. 5A), suggesting a similar or concerted evolutionary trajectory among these proteins. To precisely determine the structural localization of these adaptive changes, we used Alfaphold2^36,37^ to model the 3D protein structures of CCL20, CXCL8, CXCL10, and CXCL16 (the four most rapidly evolving chemokines in our dataset), and mapped the positively selected codons on the models. First, these models indicated that bat CC and CXC proteins share the canonical conformation of the human and mouse counterparts, comprising an N-terminal helix, three β-strands connected by the 30’s and 40’s loop and a C-terminal α-helix^38^ (Fig. 5B). Second, adaptive changes were concentrated in the N-terminus and particularly in the β-strands of chemokines (Fig. 5B). Remarkably, alignment and superimposition of the 3D protein structures revealed that several codons were common targets of positive selection in chemokines (Fig. 5C), suggesting a conserved evolutionary trajectory and/or shared selective pressures. These shared adaptive sites, which were also among the fastest evolving codons, were specifically located between the first two Cysteines (CxC) and in the 40’s loop (Fig. 5C), two key determinants for binding receptors and signal modulation^39^. These patterns are typical hallmarks of repeated evolution under pathogen-induced selective pressure.

**Figure 5.**
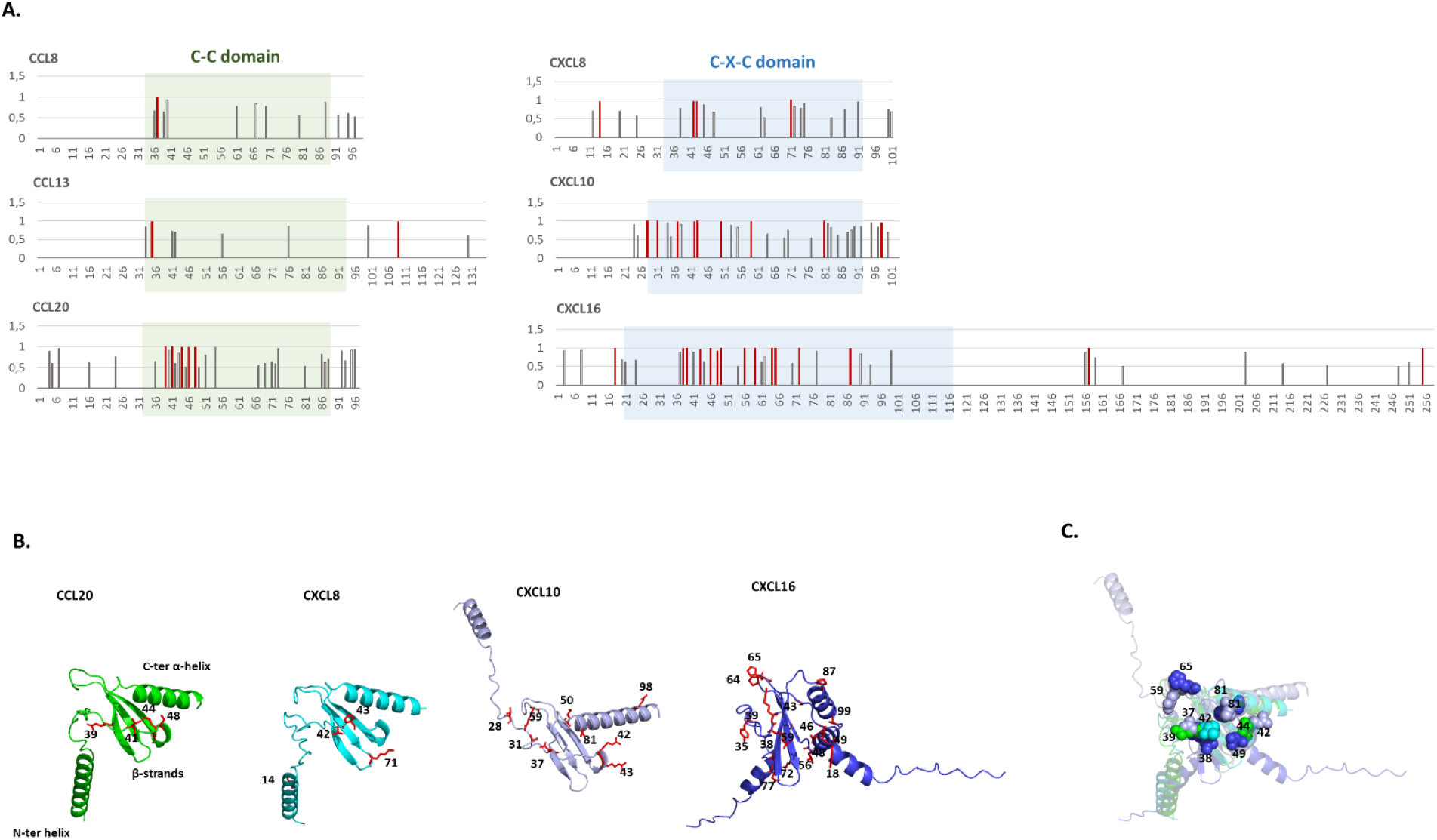
The Chemokine axis shows concerted adaptive changes targeting CC and CXC motifs. (A) Graphic representations of posterior probabilities (Bayes Empirical Bayes, BEB) from the M8 model for each codon (Y-axis). Codons under significant positive selection (BEB > 0.95) are highlighted in red. (B) Protein structure models inferred with AlphaFold 2, showing positively selected sites (in red) for chemokines with the strongest signals of positive selection. Amino acid positions correspond to the *M. myotis* protein sequence. (C) Structural alignment of chemokine proteins using the Pairwise Structure Alignment tool from the RCSB Protein Data Bank^92^. Highlighted residues correspond to *M. myotis* CXCL16.

A distinctive feature of chemokine proteins is the conserved presence of basic amino acids—arginine (R), lysine (K), and histidine (H)—in their N-terminal helix and especially within the 40’s loop^40,41^. This basic-rich region increases their binding affinity to acidic amino acids present in their receptors^40,41^. In bats, since positively selected codon hotspots were identified in the β-strands and 40’s loop of chemokines, we investigated how these evolutionary changes influenced the physicochemical properties of these regions in CCL20, CXCL8, CXCL10, and CXCL16. We found that most of the evolving codons exhibited drastic physicochemical changes, including replacement of ancestral acidic amino acids by basic or uncharged amino acids, or vice versa (e.g., CXCL16 K38 -> E, I) (Fig. 6A). Notably, several of these codons, such as A42, P43, V71 in CXCL8 and K38, T39 in CXCL16, aligned with sites previously identified as directly involved in interactions between CXC proteins and their receptors in humans^39,42^, placing them as main candidates for host-specific cytokine activity.

**Figure 6.**
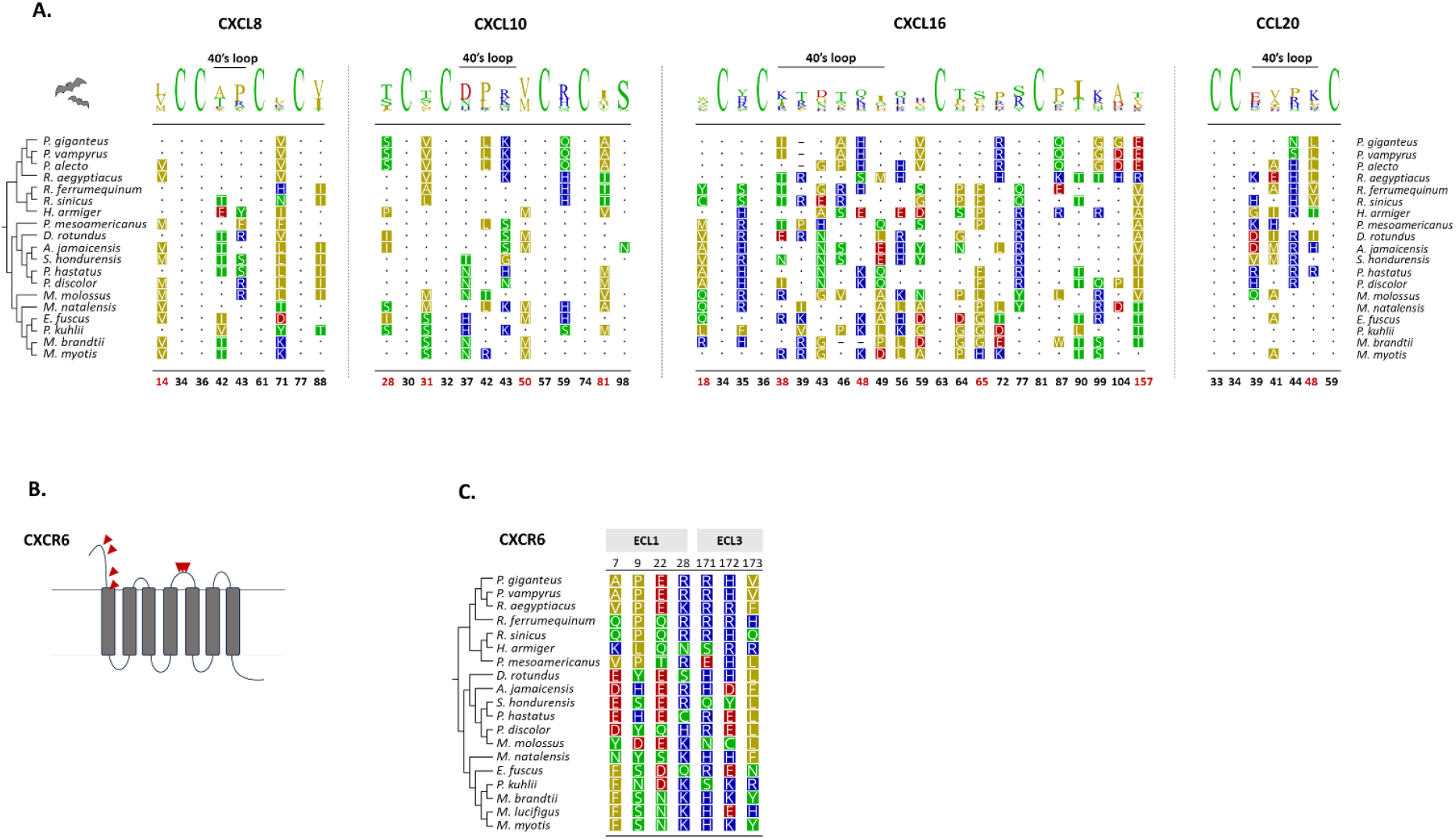
Adaptive selection of bat chemokines is associated with drastic physicochemical modifications at the receptor– ligand binding interface. (A) Alignments of positively selected sites, along conserved cysteines, highlight amino acid changes in bat chemokines. The cladogram on the left shows the phylogenetic relationships among bat species. Codons highlighted in red indicate positively selected sites identified by all five methods. The sequence logo plot generated with Geneious shows the proportion of amino acid identity at each site. (B) Schematic diagram of CXCR6 showing sites under positive selection located in the extracellular loops ECL1 and ECL3. (C) Alignments of positively selected sites in bat CXCR6 (the specific receptor of CXCL6) showing amino acid changes between species. In panels (A) and (C), numbers refer to codon positions in *M. myotis*. Colors indicate codon differences and amino acid polarity (representation from Geneious 2024.0.5).

Upon activation, chemokines first bind to glycosaminoglycans (GAGs) on the cell surface, then interact with the G-protein coupled receptors on leukocyte surfaces^40^. The rapid evolution in these binding domains may directly modulate their interaction affinity with corresponding receptors. To explore this further, we extended our evolutionary analyses to chemokine receptors, focusing on CCL20 and CXCL16, which bind to CCR3 and CXCR6, respectively. These ligand-receptor pairs offer a suitable model for investigating the evolutionary consequences of adaptive changes in chemokine binding domains. We found strong evidences of positive selection acting on CXCR6 (M7 *vs*. M8 LRT = 73, p-value = 1.9E^-16^), but not on CCR3 (M7 *vs*. M8 LRT = 6, p-value = 0.057). Notably, the positively selected codons in CXCR6 were mapped to extracellular regions, including the N-terminus (ECL1) and extracellular loop 3 (ECL3), which are critical for chemokine binding and activation^42^ (Fig. 6B). Moreover, the amino acid replacements involved radically opposite physicochemical properties, as observed in CXCL16 (Fig. 6C), consistent with adaptive plasticity at these sites. Overall, these results support a ligand-receptor co-adaptation model, highlighting that CXCL16 and CXCR6 rapidly evolved in bats.

### 4. Poxviral vCKBPs are key candidate drivers of chemokine evolution

The observed patterns of concerted evolution and co-adaptation in bat chemokines suggest that viral pressures may have played a role during chiropteran diversification. Several viruses, particularly poxviruses and herpesviruses, encode viral chemokine-binding proteins (vCKBPs) that interfere with host immunity by sequestering chemokines or antagonizing their receptors^43,44^. We therefore hypothesize that vCKBPs may act as drivers of chemokine evolution in bats. To explore this hypothesis, we focused on poxviruses, which exhibit frequent host proteins co-option events to evade or counter the immune response^45,46^. We first determined whether bat poxviruses encode vCKBPs, through Blastn search in NCBI virus database (https://www.ncbi.nlm.nih.gov/labs/virus/vssi/#/), and found homologues of these proteins in Eptesipoxvirus and Pteropoxvirus. Next, we analyzed the phylogenetic relationship between poxviral vCKBPs (from Avipoxviridae, Clade II, and bat poxviruses) and vertebrate chemokines (from avian and mammalian hosts). Surprisingly, all vCKBPs, except two encoded by Cotia viruses, clustered with the host chemokine CXCL16, the fastest evolving chemokine in our dataset. These results suggest two ancestral horizontal gene transfer of host chemokines, in particular the ancient co-option of a CXCL16-like gene into the genome of a common ancestor of most poxviruses (Fig. 7A). This is further supported by the phylogenetic structure within the vCKPB clade with the poxvirus branches closely matching the established relationships between poxviruses^47,48^. Notably, Avipoxvirus species form a distinct group, while Clade II poxviruses also show a clear clustering, reinforcing their evolutionary relationship. However, it remains unclear whether vCKBP in Avipoxvirus and Clade II poxvirus are true orthologues. Comparing the amino acid sequences of chemokines, including positively selected sites within the 40’s loop, with those of poxviral vCKBPs revealed two main patterns (Fig. 7B): (i) several conserved sites shared between host and viral-encoded chemokines, as well as among different viral-encoded chemokines (e.g., positions 10, 14, 35, 50 and 71); and (ii) highly divergent sites overlapping with positively selected sites in host chemokines (e.g., positions 38 and 48), consistent with a model of host-virus conflicts.

**Figure 7.**
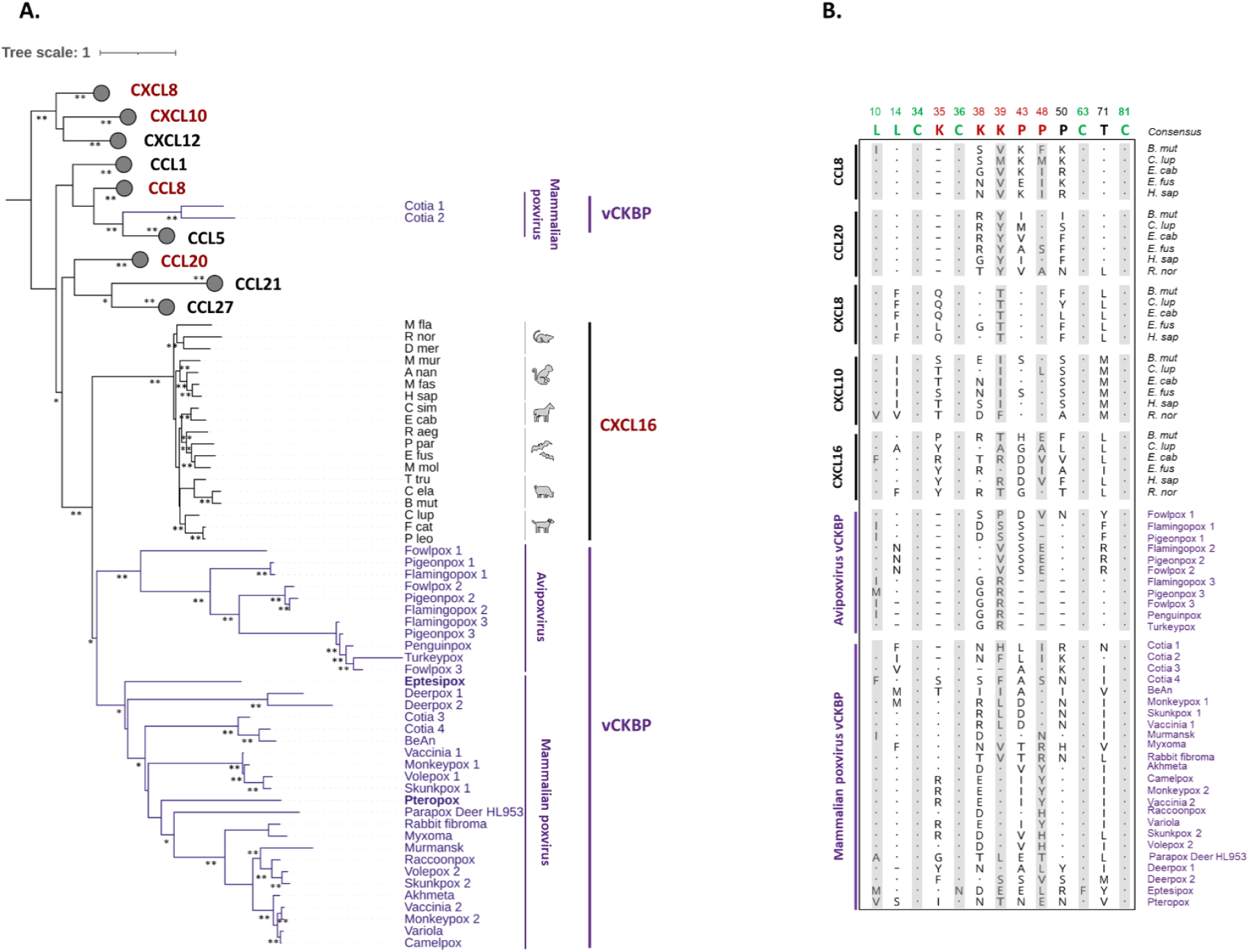
Phylogenetic analysis suggests that poxvirus vCKBP originated from at least two gene transfer events, including one involving an ancestral CXCL16-like gene. (A) Maximum-likelihood phylogenetic tree showing the relationships between vertebrate chemokines and poxviral vCKBPs. The scale bar indicates the number of substitutions per site. Bootstrap values ≥ 0.7 are indicated by asterisks: * > 0.7, ** > 0.9. The tree was rooted according to a published chemokine phylogeny^*86*^. (B) Amino acid alignment of mammalian chemokines and vCKBPs showing conserved amino acids between chemokines and vCKBPs including cysteines (green), positively selected sites in chemokines (red), and conserved amino acids among vCKBPs. Amino acid positions (indicated above the alignment) are based on *M. myotis* CXCL16. Dots represent residues identical to the consensus sequence. Species abbreviation: A. nan (*Aotus nancymaae*); B. mut (*Bos mutus*); C. ela (*Cervus elaphus*); C. lup (*Canis lupus)*; C. sim (*Ceratotherium simum*); D. mer (*Dipodomys merriami*); E. cab *(Equus Caballus*); E. fus *(Eptesicus fuscus);* F. cat (*Felis catus*); H. sap *(Homo sapiens);* M. fas *(Macaca fascicularis);* M. fla (*Marmota flaviventris*); M. mur (*Microcebus murinu*s); M. mol (*Molossus molossus*); P. leo (*Panthera leo*); P. par (*Pteronotus parnelii*); R. aeg (*Rousettus aegyptiacus*); R. nor *(Rattus norvegicus); T. tru (Tursiops truncatus)*.

## Discussion

Performing a comprehensive evolutionary analysis of the ISG transcriptome, we investigated the diversification of type I IFN pathway in bats. We uncovered widespread and strong adaptation signatures enriched in both the early detection of viral components and the downstream execution of antiviral responses, including viral dsRNA sensing, PARP-mediated ADP-ribosylation, and cytokine/chemokine signaling. Remarkably, cytokine/chemokine activity showed some of the strongest signals of positive selection, with common evolutionary patterns across several chemokine members. The concentration of genetic innovations in these molecular functions places them at the forefront of bat immunity, offering new avenues for understanding viral control, immune tolerance, and the resilience of bats to infection.

### Adaptive evolution of RNA-sensing pathways

Host immune sensors face the evolutionary challenge of recognizing highly conserved pathogen-associated molecular patterns (PAMPs), such as dsRNA, while resisting rapidly evolving viral antagonists. In bats, strong positive selection was highly concentrated in the dsRNA-binding domains of key viral RNA sensors, including TLR3, RIG-I, PKR, OAS1, and ZBP1. Although these genes have also shown evidences of adaptive evolution in other mammals (e.g., primates, rodents)^30,49,50^, such selection hotspots are unusually concentrated in the RNA-binding domains of these sensors in bats. For example, TLR3 and RIG-I in most non-bat mammals show weak signs of positive selection, typically confined to transmembrane or regulatory domains rather than the ligand-binding interfaces^30–32^. In contrast, RNA sensing domains are under strong adaptive pressure in bats, suggesting functional adaptations avoiding immune evasion and/or modulating viral detection, for example by (i) shaping the affinity or specificity in dsRNA recognition, (ii) influencing the ability to detect diverse or rapidly evolving viral dsRNA, and/or (iii) optimizing the balance in the recognition of self and non-self RNAs, ultimately maintaining immune competence while reducing the risk of autoimmune activation. This hypothesis is consistent with functional studies in humans showing that even single amino acid substitutions in the RIG-I helicase and C-terminal domains can significantly impair viral RNA recognition^51^. The recurrence of positive selection in these domains in bats further raises the possibility that bat borne viruses may have adapted to target the RNA binding regions for antagonism. However, this evolutionary dynamic remains largely unexplored and warrants further functional investigation.

### PARP gene family as an adaptive interface

We also found that members of the PARP family, particularly their macro and catalytic domains, were frequent targets of positive selection in bats. These domains are critical for ADP-ribosylation activity, a post-translational modification involved in diverse cellular processes, including DNA repair, immune regulation and antiviral defense^34,52^. In primates, PARP macro-domains also show recurrent positive selection^53,54^, highlighting their role as critical interfaces in host–pathogen conflicts. In bats, while the specific roles of PARPs remain to be characterized, their evolutionary conservation across mammals, combined with the strong selection signals we observed, suggest that PARPs may also play an important role in modulating bat innate immunity. In this context, the adaptive evolution of these domains could reflect intensified selection to maintain or enhance immune regulatory functions in response to viral antagonism, such as coronaviruses and alphaviruses that encode macrodomains capable of disrupting host PARP activity^55,56^. Additionally, in human cells, PARP9 and PARP14 co-regulate pro-inflammatory responses, with PARP9 enhancing STAT1-dependent signaling and IFN response, while PARP14 acts as a negative regulator of pro-inflammatory genes and STAT1 phosphorylation^57,58^. This dynamic regulation positions PARPs as key candidate effectors in maintaining the balance between antiviral immunity and immune tolerance, a function that may be particularly critical in Chiroptera, which may rely more heavily on non-inflammatory antiviral strategies compared to other mammals.

### Chemokines as evolutionary hotspots

Bats also exhibit elevated dN/dS ratios in their chemokines, reinforcing the likely central evolutionary role of these proteins play in bat immune system at the intersection of antiviral resistance and inflammation control. As critical mediators of immune signaling, cytokines are key players in innate immunity and inflammation, by coordinating the spatial and temporal dynamics of immune cell recruitment and activation. Maintaining this balance is crucial, as dysregulated cytokine and chemokine activity can lead to severe immunopathologies, including cytokine storms, chronic inflammation, and tissue damage^59^. Our data reveal that several positively selected sites are shared across chemokine genes, and clustered within the CC and CXC loop regions – which are critical structural motifs for receptor binding and signaling modulation. These patterns suggest either concerted evolution driven by common selective pressures or structural constraints limiting adaptation to functionally permissible regions. Rather than enhancing immune activation as a primary effect, we suggest that these adaptive changes may represent functional changes modulating the timing or magnitude of chemokine activity while escaping viral selective pressure in bats. In particular, CXCL10 was one of the most upregulated and rapidly evolving genes in our dataset, showing strong signatures of positive selection uniquely in bats. In other mammals, CXCL10 is increasingly considered as a key player in RNA and DNA virus infections^60,61^. In bats, its evolutionary pattern suggests that it may be a major target of viral selective pressure. Finally, the evolutionary footprints observed in both chemokines and their corresponding receptors, particularly in the CXCL16–CXCR6 axis, suggest ligand-receptor co-evolution driven by pathogen pressure, such as viral mechanisms to subvert, mimic, or disrupt host chemokine signaling.

### Poxvirus-encoded chemokines may be key drivers of chemokine evolution in bats

Several viruses, particularly large DNA viruses like poxviruses and herpesviruses, have evolved diverse strategies to modulate the host chemokine system. These include the production of viral chemokine homologues, viral receptors that mimic host receptors, and viral chemokine-binding proteins (vCKBPs) that act as decoy receptors to sequester host chemokines and disrupt immune signaling^43,44^. The observed adaptive changes in bat chemokines and other cytokines, such as TNFα, TNFRSF14, and TNFRSF13B, may reflect ancient arms races with herpes- and/or poxvirus-related pathogens, or other viruses that use similar antagonism mechanisms and circulated naturally in bat populations^e.g. 62,63^. Among these, poxviruses are particularly notable, as they are among the few modern viral pathogens known to cause overt disease and pathology in some bat species^62,64^. Furthermore, poxviruses have also been identified as major drivers of the adaptive evolution of bat antiviral effectors such as PKR, with clear consequences for host–virus specificity^15^. Here, we found that most poxviral vCKBPs, including those of bat-borne poxvirus, share a common ancestor with the CXCL16 clade, one of the most rapidly evolving chemokines in this dataset, thereby providing insight into the phylogenetic origin of poxviral-encoded vCKBPs. It is plausible that ancient poxviruses co-opted CXCL16-like protein homologs, possibly as decoy molecules to subvert the CXCL16–CXCR6 signaling axis. More broadly, such molecular mimicry may have exerted selective pressure on the chemokine network, contributing to its rapid evolution. Nonetheless, our results do not rule out the influence of other selective pressures. Chemokines are involved in a wide range of physiological processes, as well as interactions with other pathogens such as bacteria that encode immune inhibitors^65,66^, which are increasingly recognized as key players in host–pathogen co-evolutionary dynamics^67,68^. Further comparative functional studies are needed to clarify their roles in bats, identify any specific features compared to other mammals, and disentangle the complex interplay of selective pressures shaping their evolution.

### Broader implications beyond antiviral defense

Beyond infectious diseases, the evolution of these immune mediators may have broader implications for other physiological processes. Indeed, cytokines and chemokines not only play central roles in antiviral defense but in regulating autoimmunity, tissue repair, and tumor surveillance. Previous studies have shown many genes in the DNA repair pathway are under positive selection in bats, likely reflecting adaptations minimizing the detrimental effects of DNA damage and maintaining genome integrity^9,69^. Moreover, growing evidence suggests that bats have evolved specific mechanisms for managing cancer risk, potentially contributing to their remarkable longevity and resilience^69–71^. In our analysis, we found strong enrichment for genes associated with immune-mediated and inflammatory diseases, including autoimmune disorders, hepatitis-related pathways, and cancer-related signaling. CXCL10 and CXCL16 are known to modulate immune cell trafficking in contexts ranging from viral infections to chronic inflammation and tumor immunity^72–74^. Likewise, IL-15 plays a critical role in enhancing antitumor immunity by promoting the activation of NK and CD8+ T cells. These molecules could therefore play important roles in immune regulation and disease resistance in bats. Nevertheless, since our enrichment analyses were mostly performed on human-based databases, we acknowledge that the corresponding pathways may differ in bats. Understanding how these changes modulate immune function could offer new insights into viral clearance, inflammation regulation and immune tolerance in bats.

### Conclusion and perspectives

Our study highlights key adaptive changes in dsRNA sensors, PARPs, and cytokine/chemokine pathways, revealing critical evolutionary pathways in bats that balance antiviral defense with immune tolerance. This evolutionary framework provides valuable insights into the plasticity of the innate immune system and may inform a broader understanding of host immunity in mammals. Future work should focus on linking these genetic signatures to function and to further investigate the role of viruses in driving these adaptations.

## I. Materials and Methods

### Data Collection

To investigate the evolutionary dynamics of bat interferon-stimulated genes (ISGs), we analyzed publicly available transcriptomic data reported by Shaw et al. ^22^ (https://isg.data.cvr.ac.uk/). This study compared responses in mammalian cell lines (including those from bats, humans, brown rats, cattle, pigs, sheep, and horses) following universal type I interferon (IFN) stimulation. The dataset included two bat species: the large flying fox (*Pteropus vampyrus*) and the little brown bat (*Myotis lucifugus*). From this dataset, we selected upregulated genes with a Log_2_ fold-change (Log_2_ FC) > 2 and a false discovery rate (FDR) < 0.05 in at least one of the two bat species. The resulting gene set (Table S1) was used for further analyses of genetic and genomic adaptations across 21 available bat genomes. Coding sequences for each gene were obtained using the *M. lucifugus* Refseq protein as a query through tBLASTn searches of the “Core nucleotide” database in GenBank^75^.

Chemokine sequences orthologues were retrieved from artiodactyls, carnivores, primates, rodents, and perissodactyls, using protein sequences from *Bos mutus, Canis lupus, Homo sapiens, Mus musculus, Equus caballus* as query for tBLASTn analyses, respectively.

Representative viral chemokine-like proteins encoded by poxviruses were retrieved using annotation data from NCBI Virus database (https://www.ncbi.nlm.nih.gov/labs/virus/vssi/#/).

### Phylogenetic and positive selection analyses

Each gene dataset was codon aligned using PRANK^76^ with default parameters, except the +F option, and manually curated as necessary. Phylogenetic trees were constructed for each gene using the Maximum-Likelihood (ML) method implemented in PhyML v3.3.2^77^, with the best substitution model selected by the Smart Model Selection (SMS) program^78^.

Before assessing for positive selection, we tested for recombination events with the Genetic Algorithm for Recombination Detection (GARD)^77^. No significant recombination was detected in the genes with further analyses. Positive selection analyses were then conducted using the Branch-site Unrestricted Statistical Test for Episodic Diversification (BUSTED)^23^ from HYPHY package, as well as the PAML Codeml package^79^. The latter compares models that do not allow for positive selection (M1 and M7) with models that do (M2 and M8). These analyses included codon frequency models F61 and F3×4, with an initial omega (dN/dS ratio) of 0.4. Likelihood ratio tests were then performed to compare model pairs (M1 vs. M2 and M7 vs. M8), while codons under positive selection were identified using Bayes Empirical Bayes (BEB) analysis in M2 and M8 models, with a posterior probability threshold of ≥ 0.95 for significance. To complement these analyses, the Fast-Unbiased Bayesian Approximation (FUBAR)^80^, Fixed Effects Likelihood (FEL), and the Mixed Effects Model of Evolution (MEME)^81^, all implemented in the HYPHY package^82^, were also applied to detect positively selected codons. For higher specificity, genes and codons were considered under significant positive selection only if identified by at least two independent methods. To assess lineage-specific selective pressure in bats, we used the aBSREL model available from the Datamonkey webserver (https://www.datamonkey.org/)^83^. Given the shared evolutionary patterns observed among chemokines, gene sequences were concatenated to enhance the detection of episodic positive selection across this network using the aBSREL model.

To identify common adaptive features among chemokines and between chemokines and vCKBPs, structure-based protein alignments were performed using the PROMALS3D server^84^ (http://prodata.swmed.edu/promals3d/promals3d.php), as well as MAFFT software v.7^85^ with the L-INS-i iterative refinement method, the Dash option and default parameters. Phylogenetic relationships between host chemokines and poxviral vCKBPs were then inferred using IQ-TREE v2.4, with the best substitution model automatically detected by the software including the freeRate heterogeneity option^86^. Node statistical support was assessed through 1,000 Ultrafast bootstrap replicates. All phylogenetic trees were visualized and edited using iTOL v7, available as a free web-server tool (https://itol.embl.de/)^87^. The chemokine phylogenetic tree was rooted using CXC chemokines, following the topology established in previously published chemokine phylogenies^88^. Protein sequence logo and amino acid physico-chemical properties were visualized with Geneious Prime 2024.0.5.

### Gene Ontology and pathway enrichment analyses

To investigate the biological processes associated with genes under positive selection, we conducted functional enrichment analyses using Gene Ontology (GO) annotations and pathway databases. Specifically, GO terms were assigned and visualized, and complementary pathway information was retrieved from WikiPathways^89^ and Reactome^90^. Enrichment analyses were performed using the clusterProfiler^91^ and org.Hs.eg.db R packages. To identify representative biological processes, the enrichGO function was applied with the following parameters: pAdjstMethod = “BH”, pvalueCutoff = 0.01, and qvalueCutoff = 0.05.

### Protein structure modeling

Protein motifs and domains were identified using Interpro (https://www.ebi.ac.uk/interpro/)^92^, and diagrams with representative positively selected site were design DOG 2.0^93^. The 3D protein structures were predicted using AlphaFold2^36,37^ with default parameters. The best model was chosen based on the mean pLDDT (> 0.70) which assesses the quality of the models. RNA – Protein docking was performed via the HDOCK server^94^, this software uses a fast Fourier transform–based search strategy to model different potential binding means between the proteins, and then, each binding mode is evaluated using the scoring function ITScorePP. We used the Watson-Crick 16-mer dsRNA crystal structure (PDB 3nd4) as a ligand, and predicted protein structures of TLR3, RIGI, OAS1 and PKR RBD using *P. giganteus*’s protein sequence as a reference. We kept the top model with the highest negative score and a confidence score > 0.9. 3D protein structures of chemokines were aligned using the Pairwise Structure Alignment tool available at RCSB Protein Data Bank (RCSB PDB, https://www.rcsb.org/alignment)^95^. Inferred and superimposed protein structures were visualized and designed with PyMol v2.6.

### Additional statistical analyses

The correlation between dN/dS ratios and log_2_ fold change in gene expression was statistically assessed using Spearman’s rank correlation. Differences in average dN/dS ratios among mammalian chemokine genes were evaluated using the Kruskal-Wallis test. A P-value < 0.05 was considered statistically significant for all analyses.

### Data and code availability

All data are included within the paper and/or the Supplementary Materials. Alignments, phylogenetic trees and scripts used for evolutionary analyses are publicly available at https://gitlab.in2p3.fr/elfilali/evobats.

## Acknowledgement

This study was made possible by publicly available transcriptomic resources that adhere to FAIR principles, thereby promoting open science through data integration and reproducibility. We are grateful to the contributors of the LBBE bioinformatic server, the publicly available bioinformatic programs, and the publicly available transcriptomic and genomic data. We sincerely thank Caroline Strube for her support and helpful discussions, and acknowledge all colleagues who contributed through thoughtful discussions.

## Author contributions

Conceptualization: SJ, LE and DP. Methodology: SJ, AEF, LE, DP. Formal analysis: SJ, AEF. Visualization: SJ, AEF. Writing - original draft: SJ. Writing – review and editing: all authors. Funding: SJ, LE, DP.

## References

1. Schoggins, J. W. Interferon-Stimulated Genes: What Do They All Do? Annual Review of Virology 6, 567–584 (2019).

2. Banerjee, A. et al. Novel Insights Into Immune Systems of Bats. Front. Immunol. 11, 26 (2020).

3. Gorbunova, V., Seluanov, A. & Kennedy, B. K. The World Goes Bats: Living Longer and Tolerating Viruses. Cell Metabolism 32, 31–43 (2020).

4. Irving, A. T., Ahn, M., Goh, G., Anderson, D. E. & Wang, L.-F. Lessons from the host defences of bats, a unique viral reservoir. Nature 589, 363–370 (2021).

5. Calisher, C. H., Childs, J. E., Field, H. E., Holmes, K. V. & Schountz, T. Bats: Important reservoir hosts of emerging viruses. Clinical Microbiology Reviews 19, 531–545 (2006).

6. Brook, C. E. & Dobson, A. P. Bats as ‘special’ reservoirs for emerging zoonotic pathogens. Trends in Microbiology 23, 172–180 (2015).

7. Pavlovich, S. S. et al. The Egyptian Rousette Genome Reveals Unexpected Features of Bat Antiviral Immunity. Cell 173, 1098–1110.e18 (2018).

8. Ahn, M. et al. Dampened NLRP3-mediated inflammation in bats and implications for a special viral reservoir host. Nature Microbiology (2019) doi:10.1038/s41564-019-0371-3.

9. Zhang, G. et al. Comparative Analysis of Bat Genomes Provides Insight into the Evolution of Flight and Immunity. Science 339, 456–460 (2013).

10. Xie, J. et al. Dampened STING-Dependent Interferon Activation in Bats. Cell Host and Microbe 23, 297–301.e4 (2018).

11. Ahn, M. et al. Bat ASC2 suppresses inflammasomes and ameliorates inflammatory diseases. Cell 186, 2144–2159. e22 (2023).

12. Banerjee, A., Rapin, N., Bollinger, T. & Misra, V. Lack of inflammatory gene expression in bats: A unique role for a transcription repressor. Scientific Reports 7, (2017).

13. Zhou, P. et al. Contraction of the type I IFN locus and unusual constitutive expression of IFN-α in bats. Proceedings of the National Academy of Sciences 113, 2696–2701 (2016).

14. Hölzer, M. et al. Virus- and Interferon Alpha-Induced Transcriptomes of Cells from the Microbat Myotis daubentonii. iScience 19, 647–661 (2019).

15. Jacquet, S. et al. Adaptive duplication and genetic diversification of protein kinase R contribute to the specificity of bat-virus interactions. Science Advances 8, eadd7540 (2022).

16. Hayward, J. A. et al. Unique evolution of antiviral tetherin in bats. Journal of Virology 96, e01152–22 (2022).

17. Boys, I. N. et al. RTP4 Is a Potent IFN-Inducible Anti-flavivirus Effector Engaged in a Host-Virus Arms Race in Bats and Other Mammals. Cell Host & Microbe 28, 712–723.e9 (2020).

18. Morales, A. E. et al. Bat genomes illuminate adaptations to viral tolerance and disease resistance. Nature 638, 449–458 (2025).

19. Jebb, D. et al. Six reference-quality genomes reveal evolution of bat adaptations. Nature 583, 578–584 (2020).

20. Tian, S. et al. Comparative analyses of bat genomes identify distinct evolution of immunity in Old World fruit bats. Sci. Adv. 9, eadd0141 (2023).

21. Moreno Santillán, D. D. et al. Large-scale genome sampling reveals unique immunity and metabolic adaptations in bats. Molecular ecology 30, 6449–6467 (2021).

22. Shaw, A. E. et al. Fundamental properties of the mammalian innate immune system revealed by multispecies comparison of type I interferon responses. PLoS Biology 15, 1–23 (2017).

23. Murrell, B. et al. Gene-Wide Identification of Episodic Selection. Molecular Biology and Evolution vol. 32 1365–1371 (2015).

24. Pond, S. L. K. et al. HyPhy 2.5—A Customizable Platform for Evolutionary Hypothesis Testing Using Phylogenies. Molecular Biology and Evolution 37, 295–299 (2019).

25. Jacquet, S., Pontier, D. & Etienne, L. Rapid evolution of HERC6 and duplication of a chimeric HERC5/6 gene in rodents and bats suggest an overlooked role of HERCs in mammalian immunity. Frontiers in immunology 3232 (2020).

26. Palmer, S. N., Chappidi, S., Pinkham, C. & Hancks, D. C. Evolutionary Profile for (Host and Viral) MLKL Indicates Its Activities as a Battlefront for Extensive Counteradaptation. Molecular Biology and Evolution 38, 5405–5422 (2021).

27. Fay, E. J., Isterabadi, K., Rezanka, C. M., Le, J. & Daugherty, M. D. Evolutionary and functional analyses reveal a role for the RHIM in tuning RIPK3 activity across vertebrates. eLife 13, RP102301 (2025).

28. Mishra, S. et al. Bat RNA viruses employ viral RHIMs orchestrating species-specific cell death programs linked to Z-RNA sensing and ZBP1-RIPK3 signaling. iScience 27, (2024).

29. Le Corf, A. et al. Genomic and functional adaptations in guanylate-binding protein 5 (GBP5) highlight specificities of bat antiviral innate immunity. Preprint at 10.1101/2025.02.11.637683 (2025).

30. Areal, H., Abrantes, J. & Esteves, P. J. Signatures of positive selection in Toll-like receptor (TLR) genes in mammals. BMC Evolutionary Biology 11, 368 (2011).

31. Cagliani, R. et al. RIG-I-Like Receptors Evolved Adaptively in Mammals, with Parallel Evolution at LGP2 and RIG-I. Journal of Molecular Biology 426, 1351–1365 (2014).

32. Lemos de Matos, A., McFadden, G. & Esteves, P. J. Positive evolutionary selection on the RIG-I-like receptor genes in mammals. PLoS One 8, e81864 (2013).

33. Bai, P. Biology of poly (ADP-ribose) polymerases: the factotums of cell maintenance. Molecular cell 58, 947–958 (2015).

34. Ryan, A. P., Delgado-Rodriguez, S. E. & Daugherty, M. D. Zinc-finger PARP proteins ADP-ribosylate alphaviral proteins and are required for interferon-γ–mediated antiviral immunity. Science Advances 11, eadm6812 (2025).

35. Sokol, C. L. & Luster, A. D. The Chemokine System in Innate Immunity. Cold Spring Harb Perspect Biol 7, a016303 (2015).

36. Jumper, J. et al. Highly accurate protein structure prediction with AlphaFold. nature 596, 583–589 (2021).

37. Bryant, P., Pozzati, G. & Elofsson, A. Improved prediction of protein-protein interactions using AlphaFold2. Nature communications 13, 1265 (2022).

38. Fernandez, E. J. & Lolis, E. Structure, function, and inhibition of chemokines. Annual review of pharmacology and toxicology 42, 469–499 (2002).

39. Arimont, M. et al. Structural Analysis of Chemokine Receptor–Ligand Interactions. J. Med. Chem. 60, 4735–4779 (2017).

40. Allen, S. J., Crown, S. E. & Handel, T. M. Chemokine: receptor structure, interactions, and antagonism. Annu. Rev. Immunol. 25, 787–820 (2007).

41. Stone, M. J., Hayward, J. A., Huang, C., E. Huma, Z. & Sanchez, J. Mechanisms of regulation of the chemokine-receptor network. International journal of molecular sciences 18, 342 (2017).

42. Koenen, A. et al. The DRF motif of CXCR6 as chemokine receptor adaptation to adhesion. PLOS ONE 12, e0173486 (2017).

43. Alcami, A. Viral mimicry of cytokines, chemokines and their receptors. Nature Reviews Immunology 3, 36–50 (2003).

44. Hernaez, B. & Alcamí, A. Virus-encoded cytokine and chemokine decoy receptors. Current opinion in immunology 66, 50–56 (2020).

45. Fixsen, S. M. et al. Poxviruses capture host genes by LINE-1 retrotransposition. eLife 11, e63332 (2022).

46. Rahman, M. J. et al. LINE-1 retrotransposons facilitate horizontal gene transfer into poxviruses. eLife 11, e63327 (2022).

47. Molteni, C., Forni, D., Cagliani, R., Bravo, I. G. & Sironi, M. Evolution and diversity of nucleotide and dinucleotide composition in poxviruses. Journal of General Virology, vol. 104 (2023).

48. Yu, Z. et al. Genomic analysis of Poxviridae and exploring qualified gene sequences for phylogenetics. Comput Struct Biotechnol J 19, 5479–5486 (2021).

49. Hancks, D. C., Hartley, M. K., Hagan, C., Clark, N. L. & Elde, N. C. Overlapping patterns of rapid evolution in the nucleic acid sensors cGAS and OAS1 suggest a common mechanism of pathogen antagonism and escape. PLoS genetics 11, e1005203 (2015).

50. Elde, N. C., Child, S. J., Geballe, A. P. & Malik, H. S. Protein kinase R reveals an evolutionary model for defeating viral mimicry. Nature 457, 485–489 (2009).

51. Saito, T., Owen, D. M., Jiang, F., Marcotrigiano, J. & Gale Jr., M. Innate immunity induced by composition-dependent RIG-I recognition of hepatitis C virus RNA. Nature 454, 523–527 (2008).

52. Fehr, A. R. et al. The impact of PARPs and ADP-ribosylation on inflammation and host-pathogen interactions. Genes Dev 34, 341–359 (2020).

53. Daugherty, M. D., Young, J. M., Kerns, J. A. & Malik, H. S. Rapid Evolution of PARP Genes Suggests a Broad Role for ADP-Ribosylation in Host-Virus Conflicts. PLoS Genetics 10, (2014).

54. Delgado-Rodriguez, S. E., Ryan, A. P. & Daugherty, M. D. Recurrent loss of macrodomain activity in host immunity and viral proteins. Pathogens 12, 674 (2023).

55. Grunewald, M. E. et al. The coronavirus macrodomain is required to prevent PARP-mediated inhibition of virus replication and enhancement of IFN expression. PLoS pathogens 15, e1007756 (2019).

56. Kuny, C. V. & Sullivan, C. S. Virus–Host Interactions and the ARTD/PARP Family of Enzymes. PLoS Pathogens 12, 1–7 (2016).

57. Iwata, H. et al. PARP9 and PARP14 cross-regulate macrophage activation via STAT1 ADP-ribosylation. Nature communications 7, 12849 (2016).

58. Parthasarathy, S. & Fehr, A. R. PARP14: a key ADP-ribosylating protein in host–virus interactions? PLoS pathogens 18, e1010535 (2022).

59. Fajgenbaum, D. C. & June, C. H. Cytokine storm. New England Journal of Medicine 383, 2255–2273 (2020).

60. Caetano, A. J. et al. Spatially resolved transcriptomics reveals pro-inflammatory fibroblast involved in lymphocyte recruitment through CXCL8 and CXCL10. Elife 12, e81525 (2023).

61. Ozga, A. J. et al. CXCL10 chemokine regulates heterogeneity of the CD8+ T cell response and viral set point during chronic infection. Immunity 55, 82–97. e8 (2022).

62. David, D. et al. A novel poxvirus isolated from an Egyptian fruit bat in Israel. Veterinary Medicine & Sci 6, 587–590 (2020).

63. James, S., Donato, D., de Thoisy, B., Lavergne, A. & Lacoste, V. Novel herpesviruses in neotropical bats and their relationship with other members of the Herpesviridae family. Infection, Genetics and Evolution 84, 104367 (2020).

64. Lelli, D. et al. Hypsugopoxvirus: A Novel Poxvirus Isolated from Hypsugo savii in Italy. 1–11 (2019).

65. Crawford, M. A. et al. Identification of the bacterial protein FtsX as a unique target of chemokine-mediated antimicrobial activity against Bacillus anthracis. Proceedings of the National Academy of Sciences 108, 17159–17164 (2011).

66. Kurupati, P. et al. Chemokine-cleaving Streptococcus pyogenes protease SpyCEP is necessary and sufficient for bacterial dissemination within soft tissues and the respiratory tract. Molecular Microbiology 76, 1387–1397 (2010).

67. Carey, C. M., Apple, S. E., Hilbert, Z. A., Kay, M. S. & Elde, N. C. Diarrheal pathogens trigger rapid evolution of the guanylate cyclase-C signaling axis in bats. Cell Host & Microbe 29, 1342–1350.e5 (2021).

68. Hilbert, Z. A. et al. Rapid Evolution of Glycan Recognition Receptors Reveals an Axis of Host–Microbe Arms Races beyond Canonical Protein–Protein Interfaces. Genome Biology and Evolution 15, evad119 (2023).

69. Vazquez, J. M. et al. Extensive longevity and DNA virus-driven adaptation in nearctic Myotis bats. Preprint at 10.1101/2024.10.10.617725 (2024).

70. Scheben, A. et al. Long-read sequencing reveals rapid evolution of immunity-and cancer-related genes in bats. Genome biology and evolution 15, evad148 (2023).

71. Hua, R. et al. Experimental evidence for cancer resistance in a bat species. Nature Communications 15, 1401 (2024).

72. Karin, N. & Razon, H. Chemokines beyond chemo-attraction: CXCL10 and its significant role in cancer and autoimmunity. Cytokine 109, 24–28 (2018).

73. Lee, E. Y., Lee, Z.-H. & Song, Y. W. CXCL10 and autoimmune diseases. Autoimmunity Reviews 8, 379–383 (2009).

74. Bao, N. et al. Role of the CXCR6/CXCL16 axis in autoimmune diseases. International Immunopharmacology 121, 110530 (2023).

75. Sayers, E. W. et al. GenBank. Nucleic Acids Research 47, D94–D99 (2018).

76. Löytynoja, A. Phylogeny-aware alignment with PRANK. Methods Mol Biol 1079, 155–170 (2014).

77. Guindon, S. et al. New algorithms and methods to estimate maximum-likelihood phylogenies: Assessing the performance of PhyML 3.0. Systematic Biology 59, 307–321 (2010).

78. Lefort, V., Longueville, J. E. & Gascuel, O. SMS: Smart Model Selection in PhyML. Molecular biology and evolution 34, 2422–2424 (2017).

79. Yang, Z. PAML 4: phylogenetic analysis by maximum likelihood. Molecular biology and evolution 24, 1586–1591 (2007).

80. Murrell, B. et al. FUBAR: a fast, unconstrained bayesian approximation for inferring selection. Molecular biology and evolution 30, 1196–1205 (2013).

81. Murrell, B. et al. Detecting Individual Sites Subject to Episodic Diversifying Selection. PLOS Genetics 8, e1002764 (2012).

82. Pond, S. L. K., Frost, S. D. W. & Muse, S. V. HyPhy: hypothesis testing using phylogenies. Bioinformatics 21, 676–679 (2004).

83. Weaver, S. et al. Datamonkey 2.0: A modern web application for characterizing selective and other evolutionary processes. Molecular Biology and Evolution 35, 773–777 (2018).

84. Pei, J., Tang, M. & Grishin, N. V. PROMALS3D web server for accurate multiple protein sequence and structure alignments. Nucleic Acids Res 36, W30–34 (2008).

85. Katoh, K. & Standley, D. M. MAFFT multiple sequence alignment software version 7: improvements in performance and usability. Molecular biology and evolution 30, 772–780 (2013).

86. Minh, B. Q. et al. IQ-TREE 2: New Models and Efficient Methods for Phylogenetic Inference in the Genomic Era. Molecular Biology and Evolution vol. 37 1530–1534 (2020).

87. Letunic, I. & Bork, P. Interactive Tree of Life (iTOL) v6: recent updates to the phylogenetic tree display and annotation tool. Nucleic Acids Research vol. 52 W78–W82 (2024).

88. Aleotti, A., Goulty, M., Lewis, C., Giorgini, F. & Feuda, R. The origin, evolution, and molecular diversity of the chemokine system. Life Science Alliance 7, (2024).

89. Agrawal, A. et al. WikiPathways 2024: next generation pathway database. Nucleic acids research 52, D679–D689 (2024).

90. Milacic, M. et al. The reactome pathway knowledgebase 2024. Nucleic acids research 52, D672–D678 (2024).

91. Yu, G., Wang, L.-G., Han, Y. & He, Q.-Y. clusterProfiler: an R package for comparing biological themes among gene clusters. Omics: a journal of integrative biology 16, 284–287 (2012).

92. Blum, M. et al. InterPro: the protein sequence classification resource in 2025. Nucleic Acids Research vol. 53 D444–D456 (2024).

93. Ren, J. et al. DOG 1.0: illustrator of protein domain structures. Cell Research 19, 271–273 (2009).

94. Yan, Y., Tao, H., He, J. & Huang, S.-Y. The HDOCK server for integrated protein–protein docking. Nature protocols 15, 1829–1852 (2020).

95. Bittrich, S., Segura, J., Duarte, J. M., Burley, S. K. & Rose, Y. RCSB protein Data Bank: exploring protein 3D similarities via comprehensive structural alignments. Bioinformatics 40, btae370 (2024).

