## Supplementary Info for "Extensive adaptive changes in bat interferon pathway reveal specific molecular functions at the forefront of host–virus coevolution"

### Supplementary Information

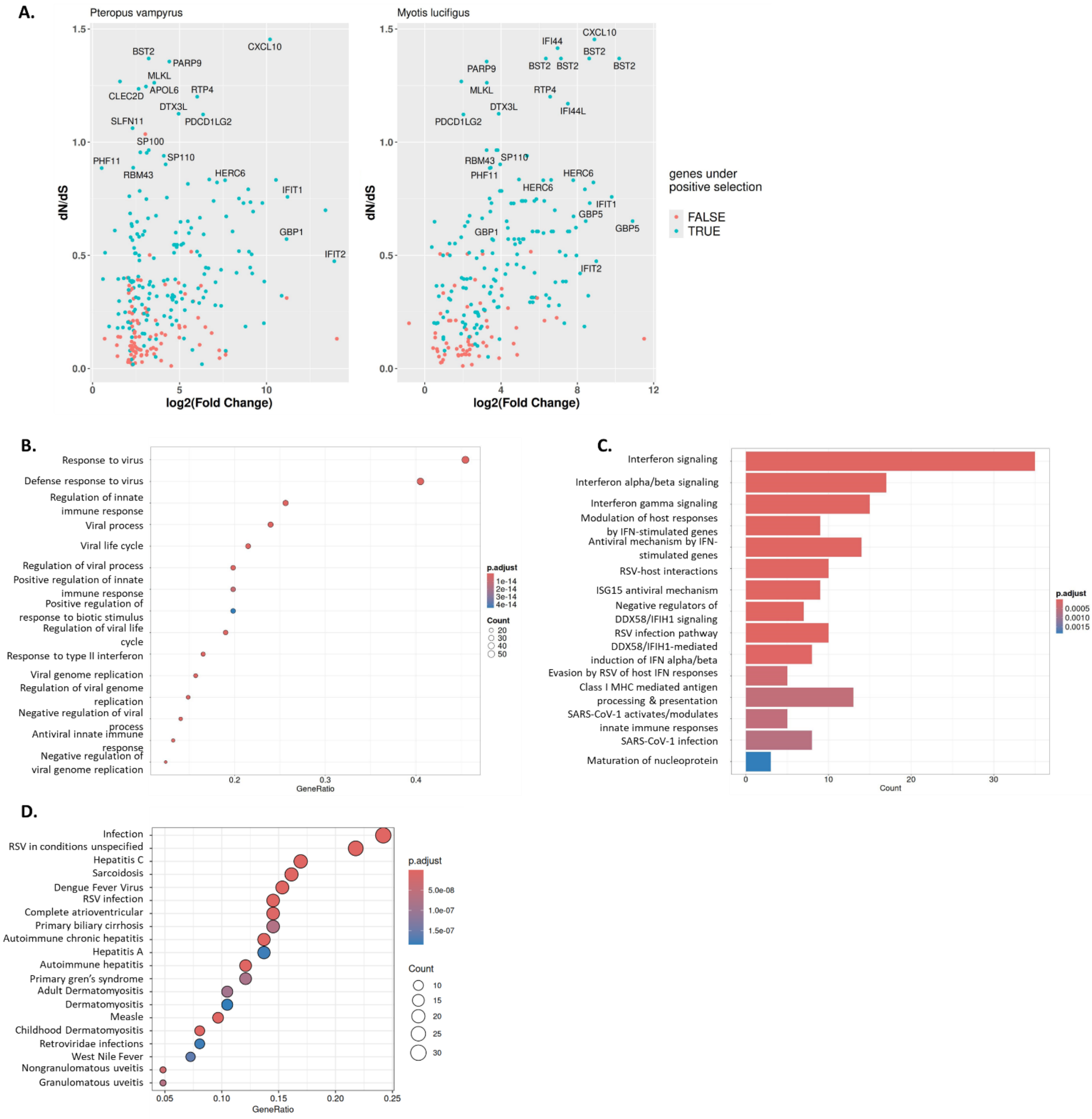


**Supplementary Figure 1 (related to Figures 1 and 2). Positive selection at the interface between bat ISGs and viral infections.** (A) Average dN/dS values plotted against $\log_2$(fold change) in *P. vampyrus* and *M. lucifugus*, highlighting ISGs under significant positive selection as identified by at least one evolutionary model (blue dots; p-value < 0.05). (B-D) Gene Ontology (GO) and DisGeNET enrichment analyses showing the biological processes, pathways, and disease association linked to the positively selected genes.

**A.**

| | M1 vs M2 | | M7 vs M8 | | BUSTED | Positively selected sites |
|---|---|---|---|---|---|---|
| **Genes** | **-2ΔLn L** | **p-value** | **-2ΔLn L** | **p-value** | **p-value** | |
| **IL15** | 23,428 | 0,000 | 28,368 | 0,000 | 0,225 | 59, 99, 112, 115, 116, 120, 152 |
| **TNF** | 15,236 | 0,000 | 34,501 | 0,000 | 0,500 | 123, 125, 133, 170 |
| **TNFSF10** | 28,836 | 0,000 | 36,550 | 0,000 | 0,118 | 6, 14, 109, 111, 112, 128, 133, 134, 135 |
| **TNFSF13B** | 8,136 | 0,017 | 8,063 | 0,018 | 0,500 | 27, 140, 277 |

**B.**

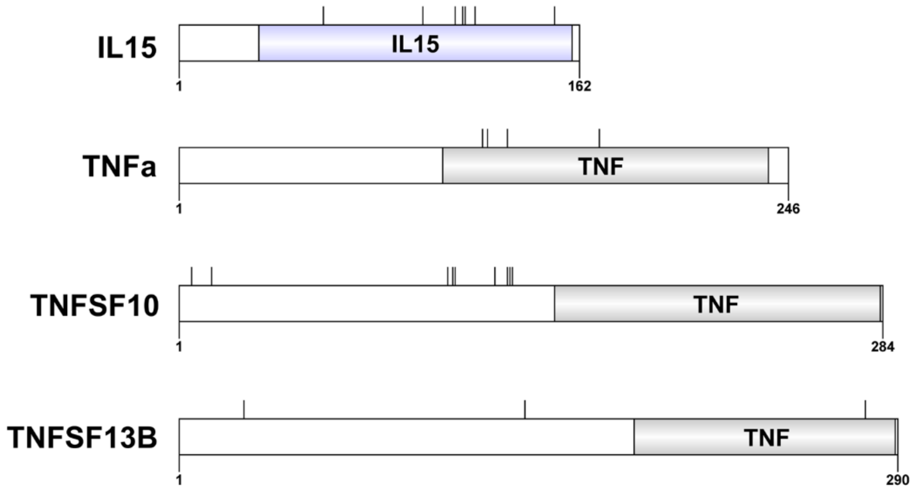


**Supplementary Figure 2. Adaptive selection in bats cytokines.** (A) Results of positive selection analyses for IL-15 and members of the TNF superfamily. (B) 2D protein domain diagrams of IL-15 and TNF superfamily members, showing positively selected sites (BEB > 0.95 or p-value < 0.05).

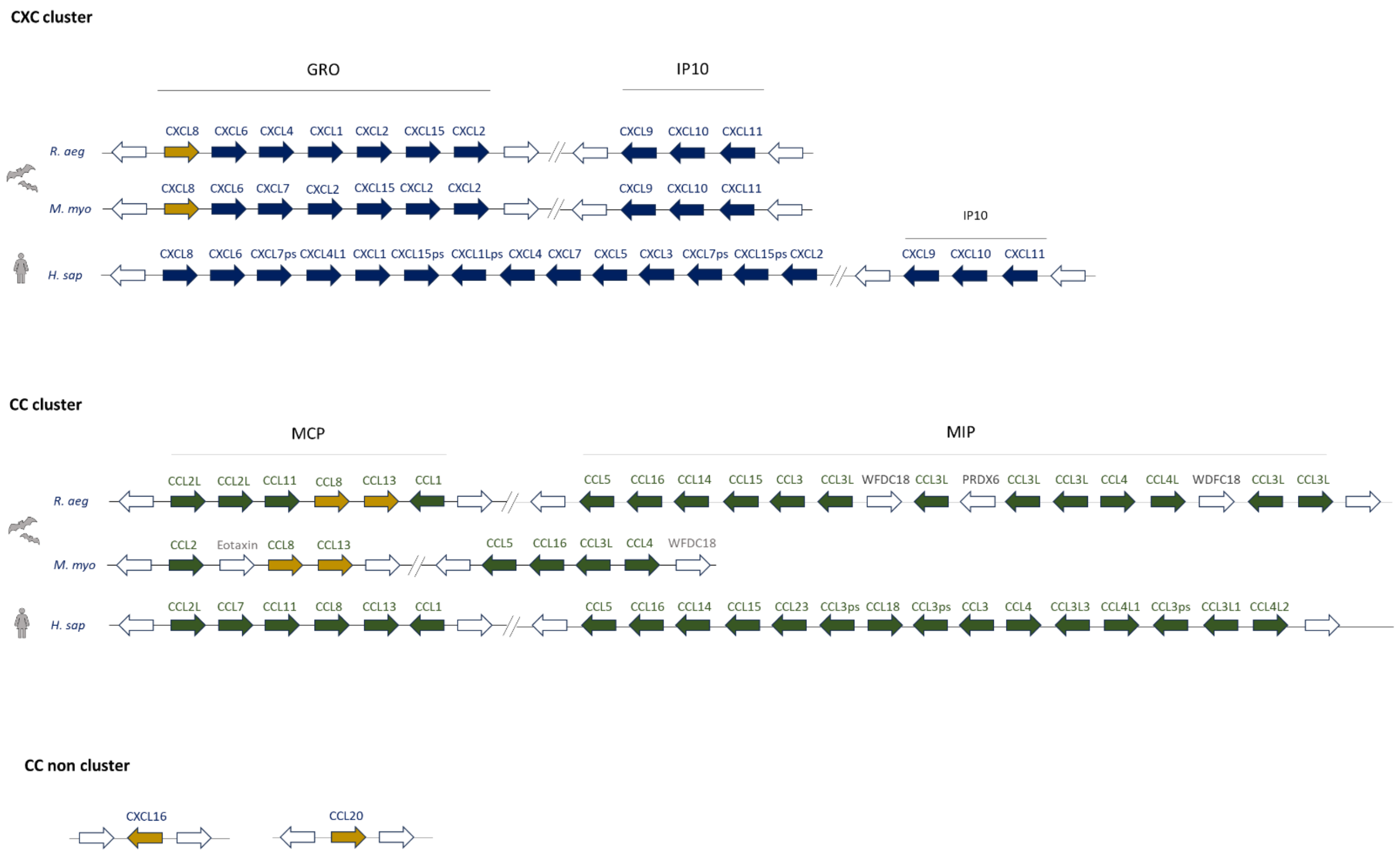


**Supplementary Figure 3 (related to Figure 4 and 5).** Genomic organizations of CXC and CC chemokine gene clusters in bat genomes from two species (*R. aeg*: *R. aegyptiacus* and *M. myo*: *M. myotis*) compared to the human genome. Gene order and orientation are shown for each locus.

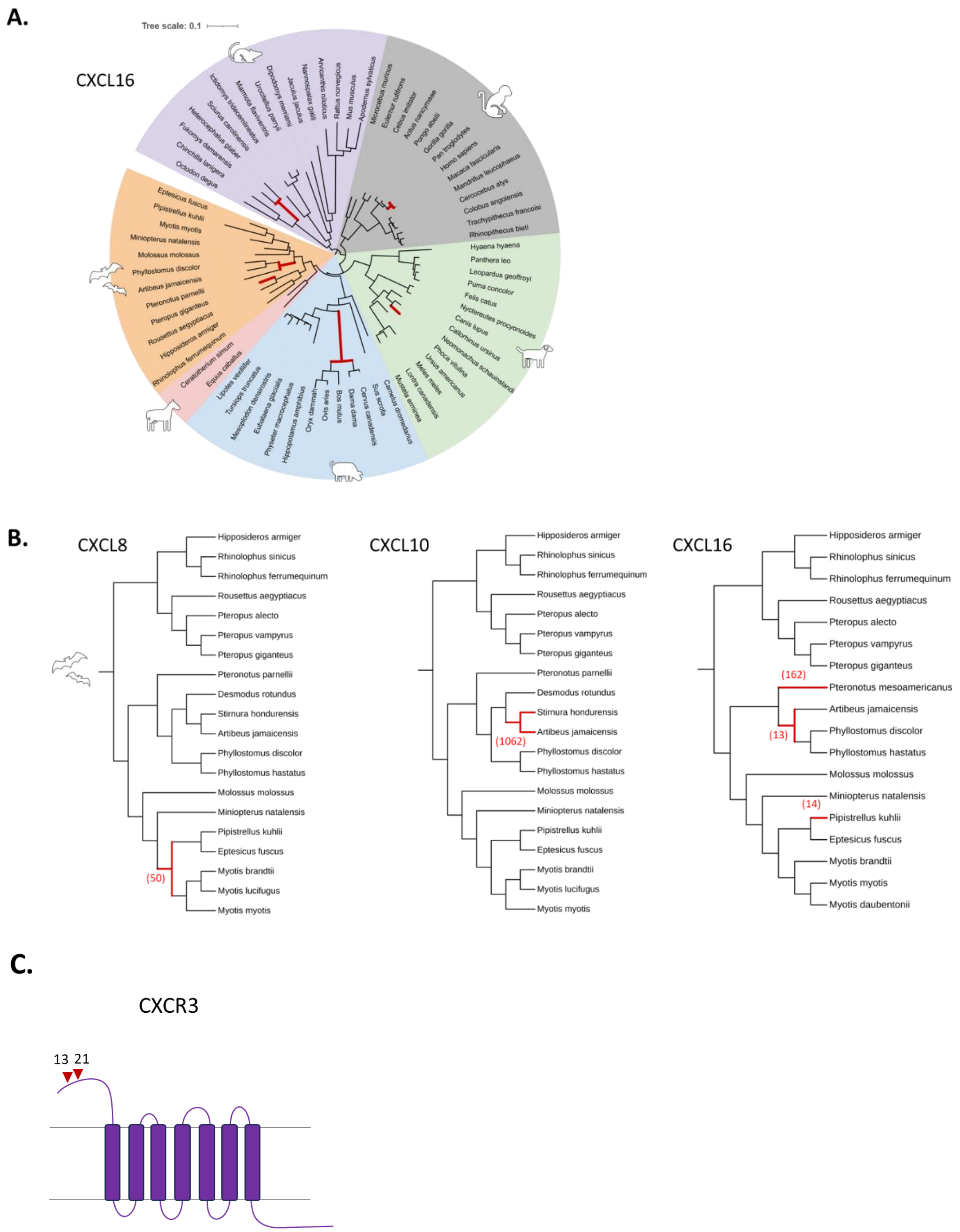


**Supplementary Figure 4 (related to Figure 3).** (A) Maximum likelihood phylogenetic trees of mammalian CCL20, CXCL16, respectively. (B) Phylogenetic tree of bat chemokine genes, showing branches under significant positive selection (p-value <0.05, in red). Numbers in brackets indicate estimated ω (dN/dS) value for each branch. The scale bar indicates the proportion of genetic variation. (C) Schematic diagram showing sites under positive selection located in the extracellular loops ECL1 of CXCR3 (the receptor of CXCL10). Amino acid positions refer to the *M. myotis* sequence.
